# An extracellular vesicle targeting ligand that binds to Arc proteins and facilitates Arc transport *in vivo*

**DOI:** 10.1101/2022.09.06.506798

**Authors:** Peter H. Lee, Michael Anaya, Mark S. Ladinsky, Justin M. Reitsma, Kai Zinn

## Abstract

Communication between distant cells can be mediated by extracellular vesicles (EVs) that deliver proteins and RNAs to recipient cells. Little is known about how EVs are targeted to specific cell types. Here we identify the *Drosophila* cell-surface protein Stranded at second (Sas) as a targeting ligand for EVs. Full-length Sas is present in EV preparations from transfected *Drosophila* Schneider 2 (S2 cells). Sas is a binding partner for the Ptp10D receptor tyrosine phosphatase, and Sas-bearing EVs preferentially target to cells expressing Ptp10D. We used co-immunoprecipitation and peptide binding to show that the cytoplasmic domain (ICD) of Sas binds to dArc1. dArc1 and mammalian Arc are related to retrotransposon Gag proteins. They form virus-like capsids which encapsulate *Arc* and other mRNAs and are transported between cells via EVs. The Sas ICD contains a motif required for dArc1 binding that is shared by the mammalian and *Drosophila* amyloid precursor protein (APP) orthologs, and the Sas and APP ICDs also bind to mammalian Arc. Sas facilitates delivery of dArc1 capsids bearing *dArc1* mRNA into distant Ptp10D-expressing recipient cells *in vivo*.

## INTRODUCTION

Extracellular vesicles (EVs) are mediators of cell-cell communication that transport specific protein and RNA cargoes. They are a heterogeneous collection of vesicular structures that are exported from cells by a variety of mechanisms. Exosomes are 30-150 nm in diameter and are released into cell supernatants *via* fusion of multivesicular bodies (MVBs) with the plasma membrane. Exosomes and other EVs carry specific proteins and RNAs, and EVs derived from different cell types contain different cargoes. EV cargoes are biomarkers for specific diseases. Because EVs can encapsulate RNAs and protect them from degradation, and then deliver those RNAs to recipient cells, they represent a promising new type of therapeutic agent(O’Brien et al., 2020; Teng and Fussenegger, 2020).

While the biogenesis of EVs is comparatively well understood, much less is known about mechanisms involved in their targeting to specific cell types. EVs can directly activate intracellular signaling by interacting with cell surface receptors. They are internalized into cells after receptor binding using a variety of endocytic mechanisms, resulting in the delivery of their cargoes into the recipient cells. In this paper, we identify Stranded at second (Sas), a large *Drosophila* cell surface protein (CSP)(Schonbaum et al., 1992), as an EV targeting ligand. Sas has an extracellular domain (ECD) containing a signal peptide, a unique N-terminal region, four von Willebrand factor C (VWFC) domains, and three Fibronectin Type III (FN-III) repeats (Fig. 1a). It has a single transmembrane (TM) domain and a short (37 amino acids (aa)) cytoplasmic domain (ICD). Sas is commonly used as a marker for the apical surfaces of epithelially-derived cells, including tracheal cells in the respiratory system. *sas* mutant larvae die at or before second instar (hence the name stranded at second, which is derived from baseball terminology) and have tracheal phenotypes(Schonbaum *et al*., 1992). A tyrosine motif in the Sas ICD binds to the PTB domain of Numb(Chien et al., 1998), an endocytic protein that is a negative regulator of Notch. Sas has no mammalian orthologs, but there are many mammalian CSPs that contain VWFC and FN-III domains.

**Fig. 1.**
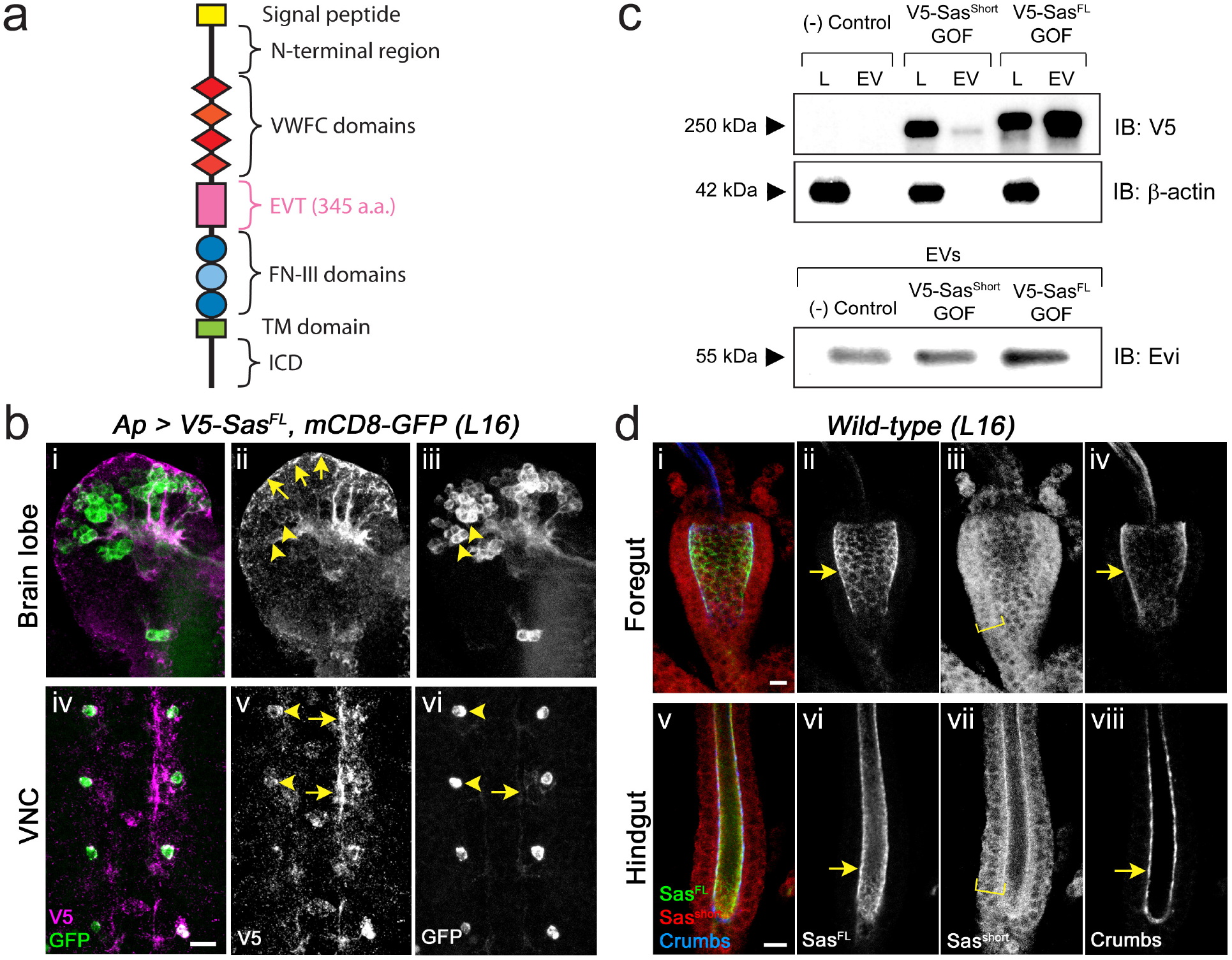
Localization of Sas isoforms. **a**, schematic diagram of the Sas^FL^ protein. The Sas^short^ isoform lacks the EVT region. **b**, Sas^FL^ moves away from expressing cells. V5-Sas^FL^ was expressed together with mCD8-GFP (transmembrane CSP) in Apterous neurons in late stage 16 embryos. **i**,**iv**: double-labeling (V5, magenta; GFP, green); **ii, v**: V5 channel; **iii, vi**: GFP channel. **i-iii**, brain lobes. GFP labels cell bodies (arrowheads in **iii**) and axon tracts. V5 labels cell bodies only weakly (arrowheads in **ii**), strongly labels some axon tracts, and localizes to the periphery (sheath) of the brain lobes (arrows in **ii**). **iv-vi**, ventral nerve cord. GFP strongly labels Ap VNC cell bodies (arrowheads in **vi**) and weakly labels Ap axons (arrow in **vi**). V5 weakly labels cell bodies (arrowheads in **v**), and strongly labels segments of axons (arrows in **v**). Note that V5 staining appears thicker than GFP staining, suggesting that it represents glial sheaths surrounding the axon tracts. Scale bar, 10 μm. **c**, Western blot, showing that Sas^FL^ localizes to EVs. EVs were prepared from cell supernatants, and equal amounts of cell lysate proteins and EV proteins were loaded on the gel. GOF (gain of function): the indicated protein is overexpressed. Top panel, anti-V5 blot. Sas^FL^ migrates slightly above the 250 kD marker, and Sas^short^ slightly below it. Middle panel, anti-β-actin (cytoplasmic marker) blot. Note that there is more Sas^FL^ in EVs than in the lysate, while almost all Sas^short^ is in lysate. The absence of β-actin signal in the EVs shows that they are not heavily contaminated by cytosol. Bottom panel, Evi (EV marker), in EV preps. **d**, Localization of endogenous Sas isoforms in the embryonic gut. *Wild-type* late stage 16 embryos were triple-stained for Sas^FL^ (using the (Schonbaum *et al*., 1992) antiserum, which primarily recognizes the EVT region; green), Sas^short^ (using our antipeptide antibody; red), and Crumbs (apical marker; blue). **i-iv**, foregut; **v-viii**, hindgut. Note that in both gut regions Sas^FL^ colocalizes with Crumbs at the apical (luminal) cell surfaces (arrows), while anti-Sas^short^ labels the entire width of the gut wall (brackets). See Supp. Fig. 1 for images of anti-Sas^short^ staining of wild-type and *sas* mutant embryos, demonstrating antibody specificity. Scale bar in **d-i**, 20 μm; in **d-v**, 10 μm. Source data files include raw and labelled images for the Western blots shown in panel c.

We identified the receptor tyrosine phosphatase (RPTP) Ptp10D as a binding partner for Sas, and showed that Sas::Ptp10D interactions regulate embryonic axon guidance, as well as glial migration and proliferation(Lee et al., 2013). Ptp10D is one of the two *Drosophila* R3 subfamily RPTPs, which have ECDs composed of long chains of FN-III repeats. Sas::Ptp10D interactions also control the elimination of neoplastic epithelial clones by surrounding normal tissue. Sas is on normal epithelial cells, and it relocalizes to the parts of their cell surfaces that are adjacent to the neoplastic clone and binds to Ptp10D on the neoplastic cells. Ptp10D in turn relocalizes and dephosphorylates the EGF receptor tyrosine kinase, leading to death of the neoplastic cells(Yamamoto et al., 2017). The Sas ECD probably has other binding partners as well, because it interacts with cells that do not express Ptp10D in live embryo staining assays(Lee *et al*., 2013).

Sas localizes to EVs, as demonstrated by immuno-electron microscopy (immuno-EM) and Western blotting of EV preparations. These EVs preferentially target to cells expressing Ptp10D, and expression of Numb further increases incorporation of EV contents into recipient cell lysates. We used mass spectrometry to identify proteins associated with Sas in EVs, and found that dArc1 is the most highly enriched protein. We then used co-immunoprecipitation (co-IP) and peptide binding to show that dArc1 binds directly to the short Sas ICD.

Arc was originally identified in mammals as a locally translated dendritic protein that regulates synaptic plasticity, in part by modulating endocytosis of AMPA receptors(Chowdhury et al., 2006; Shepherd et al., 2006). The *Drosophila* genome encodes two Arc-related proteins, dArc1 and dArc2. The *dArc2* gene, which encodes a truncated protein, was likely generated by a gene duplication, and the *dArc1* and *dArc2* genes are adjacent(Mattaliano et al., 2007). dArc1 functions in larval and adult brain neurons to regulate aspects of metabolism(Keith et al., 2021; Mattaliano *et al*., 2007; Mosher et al., 2015). Arc and dArc1 evolved independently from retrotransposon Gag proteins(Shepherd, 2018). Remarkably, they were both recently shown to form virus-like capsids that can encapsulate *Arc* mRNAs and are transported between cells via EVs(Ashley et al., 2018; Hantak et al., 2021; Pastuzyn et al., 2018). dArc1, but not dArc2, has a C-terminal Zn^2+^ finger that might be involved in nucleic acid binding(Erlendsson et al., 2020; Pastuzyn *et al*., 2018). Mammalian Arc lacks Zn^2+^ fingers, but RNA is required for normal capsid assembly(Pastuzyn *et al*., 2018). *Drosophila* dArc1 capsids bearing *dArc1* mRNA move from neurons to muscles across larval neuromuscular junction (NMJ) synapses, and dArc1 transfer is required for activity-induced induction of morphological synaptic plasticity(Ashley *et al*., 2018).

The short Sas ICD contains a tyrosine motif required for dArc1 binding. Appl, the ortholog of amyloid precursor protein (APP), is the only other Drosophila CSP that shares this motif, and its ICD also binds to dArc1. The motif is conserved in human APP, and the APP and Sas ICDs also bind to mammalian Arc. The interaction between APP and Arc is of interest because several studies have implicated Arc in control of β-amyloid accumulation and Alzheimer’s disease (AD)(Bi et al., 2018; Landgren et al., 2012; Wu et al., 2011), and APP also localizes to EVs(Laulagnier et al., 2018; Perez-Gonzalez et al., 2020).

To determine whether Sas can target dArc1 to Ptp10D-expressing recipient cells *in vivo*, we expressed dArc1 with and without Sas in embryonic salivary glands (SGs). We observed that expression of dArc1 protein from a cDNA construct induces expression of the endogenous *dArc1* gene in SGs. When Sas and dArc1 are expressed together in SGs, high levels of endogenous *dArc1* mRNA appear in distant tracheal cells, which express Ptp10D. The data suggest that Sas EVs bearing dArc1 capsids that contain *dArc1* mRNA travel within the embryo and are internalized into tracheal cells, which then also turn on expression of the endogenous *dArc1* gene.

## RESULTS AND DISCUSSION

### Sas is an EV targeting ligand

Sas exists as two isoforms generated by alternative splicing. Full-length Sas (PB/PD isoform, denoted here as Sas^FL^) is a 1693 aa protein. It contains a 345 aa region (EVT) between the VWFC and FN-III domains that is lacking in the PA/PC isoform (Sas^short^)(Fig. 1a). We expressed Sas^FL^ tagged with an N-terminal V5 epitope tag (inserted immediately after the signal sequence) in embryonic late stage 16 Apterous (Ap) neurons, which consist of paired neurons (one per hemisegment) in the ventral nerve cord (VNC) and scattered neurons in the brain lobes. We noted that V5-Sas^FL^ moved away from the expressing cells and accumulated in sheaths around brain lobes and around axons in the VNC, as well as in puncta throughout the VNC and brain (Fig. 1b). This was surprising, since Sas^FL^ is a transmembrane CSP. It was expressed together with mCD8-GFP, which is also a transmembrane CSP, and the GFP signal was restricted to the Ap neuron cell bodies, with faint staining on the axons (Fig. 1b).

Movement of V5-Sas^FL^, and presumably of endogenous Sas^FL^, away from its source could occur through cleavage of the Sas ECD from the cell surface or by release of intact Sas in EVs. To distinguish between these possibilities, we expressed V5-tagged Sas^FL^ and Sas^short^ in *Drosophila* Schneider 2 (S2) cells in culture, prepared EVs from cell supernatants using the Invitrogen Exosome Isolation Kit, and analyzed their contents by Western blotting. EV preparations generated with this kit have been shown to have similar characteristics to those generated by ultracentrifugation(Skottvoll et al., 2019). Both preparations contain primarily exosomal proteins. However, they also contain similar levels of proteins annotated as components of other compartments, especially nuclear proteins. Thus, both methods should be regarded as enrichments rather than purifications. The kit has the advantage of requiring much less material, making it suitable for generation of EVs from small populations of transfected cells. EVs generated from S2 cells contain the Evi protein, which is a commonly used EV marker. Evi is also present in cell lysates, however. The EV preparations lack β-actin, showing that they are not heavily contaminated by cytosol (Fig. 1c). We found that most of the V5-Sas^FL^ localized to EVs, while V5-Sas^short^ was retained in the cell lysate (Fig. 1c). We did not observe any proteolytic cleavage products in EVs or unpurified supernatants. Endogenous Sas is expressed at almost undetectable levels in S2 cells.

The commonly used rabbit antiserum against Sas primarily recognizes the EVT region, so cell staining reveals the localization of Sas^FL^. (Schonbaum *et al*., 1992) To visualize Sas^short^, we made an anti-peptide antibody against a sequence spanning an exon junction in the PA/PC isoforms. This selectively recognizes Sas^short^ (Supp. Fig. 1). Double-staining of the foregut and hindgut with the two Sas antibodies shows that Sas^FL^ localizes to apical cell surfaces, while Sas^short^ is distributed across the entire cell membrane (Fig. 1d). These data imply that the EVT sequence lacking in Sas^short^ is required for both apical localization and targeting to EVs. Polarized cells can release EVs with different cargoes from their apical and basolateral surfaces(Matsui et al., 2021), so EV targeting could be downstream of apical localization *in vivo*. S2 cells are unpolarized, however, so this mechanism is unlikely to apply to localization of Sas^FL^ to EVs in cultured S2s.

**Supp. Fig. 1.**
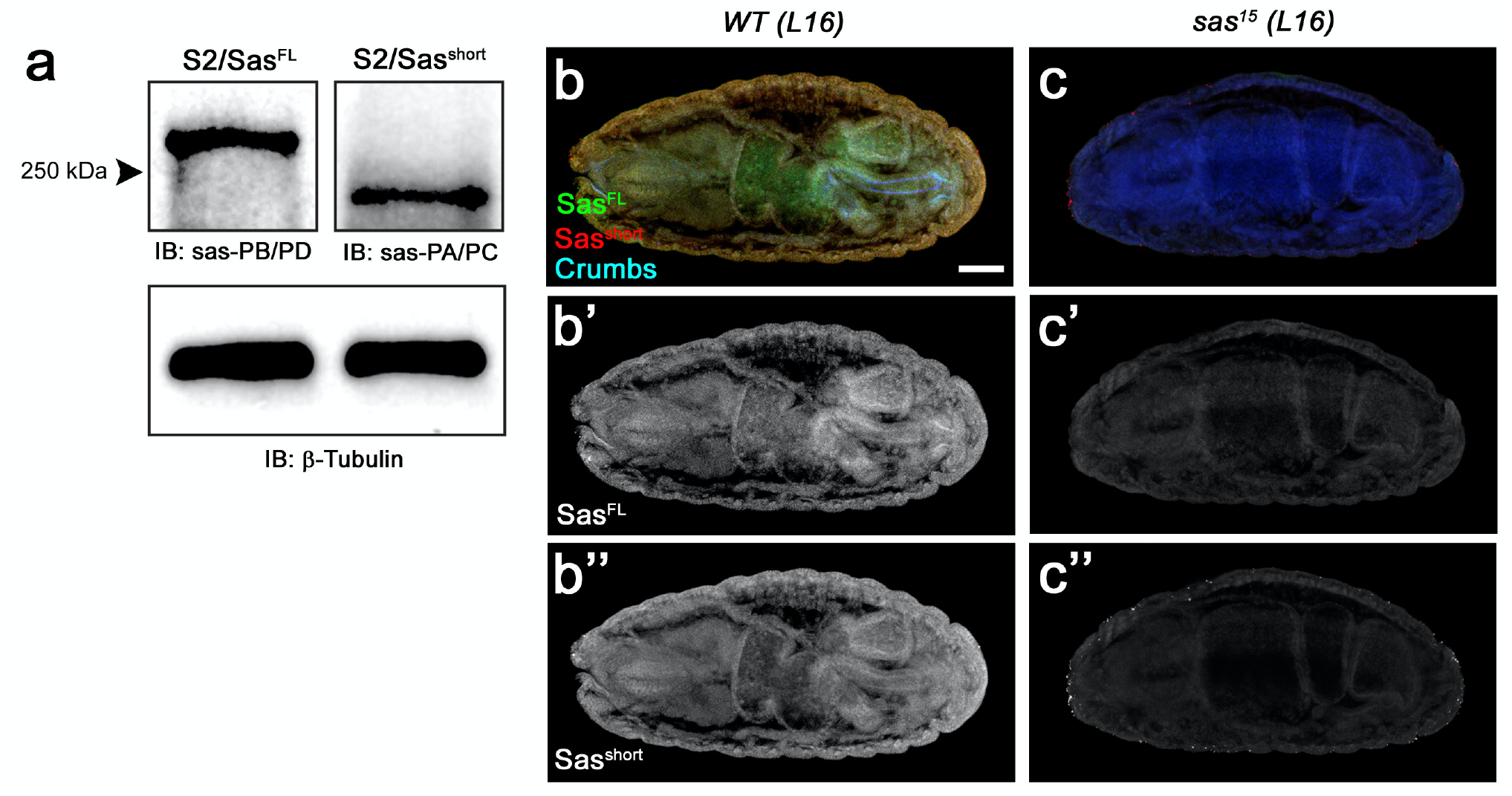
Recognition of Sas isoforms by anti-Sas antibodies. **a**, Western blot of S2 cell lysates with antibodies against Sas^FL^ (Sas-PB/PD)(Schonbaum *et al*., 1992) and Sas^short^ (Sas PA/PC) (see Materials and Methods). Left, S2 cells expressing Sas^FL^; right, S2 cells expressing Sas^short^. Anti-Sas-PB/PD recognizes a band of >250 kD, while anti-Sas-PA/PC recognizes a smaller band. **b**,**c**, late stage 16 *wild-type* (**b**) and *sas*^*15*^ (null mutant) (**c**) whole embryos, triple-stained with anti-Sas^FL^, anti-Sas^short^, and anti-Crumbs. **b’, c’** show the anti-Sas^FL^ channel only, and **b’’, c’’** show the anti-Sas^short^ channel. Note that there is no detectable staining of the *sas* mutant embryo with either antibody, showing that the anti-Sas^short^ antibody recognizes Sas *in vivo* and does not detectably cross-react with other proteins (**c’, c’’**). Scale bar, 50 μm. Source data files include raw and labelled images for the Western blots shown in panel a.

### Analysis of Sas^FL^ EVs by electron microscopy

To demonstrate that Sas is actually on EVs, we used immuno-EM and EM tomography to analyze purified EV preparations from V5-Sas^Fl^-expressing S2 cells. The tomographic images show that the EVs span a range of sizes, from ∼30 nm in diameter to >100 nm, and that they are a mixture of single and double-membrane vesicles (Fig. 2c, Supp. Figs. 2a, d). For immuno-EM, we incubated EVs with anti-V5, followed by gold-labeled anti-mouse secondary antibody. Fig. 2a shows a typical image, in which an EV is associated with multiple 10 nm gold particles. The distance between the EV membrane (yellow bracket: diameter of the vesicle) and a gold particle (white bracket: distance between membrane and a particle) varies, but can be more than 40 nm. This likely reflects the large size of the Sas ECD, in which the N-terminal V5 epitope is separated by 1590 aa from the TM domain. The region outside of the membrane boundary is of higher density, probably because it represents the protein sheath around the EV membrane. To further characterize Sas localization, we then performed an experiment in which EVs were incubated with both mouse anti-V5 and rabbit anti-Sas, which primarily recognizes the EVT region in the middle of the ECD, followed by 10 nm gold particle-labeled anti-mouse secondary antibody and 5 nm gold particle-labeled anti-rabbit secondary antibody. Fig. 2b shows an EV that is associated with multiple 10 nm (arrow) and 5 nm (arrowhead) gold particles.

**Fig. 2.**
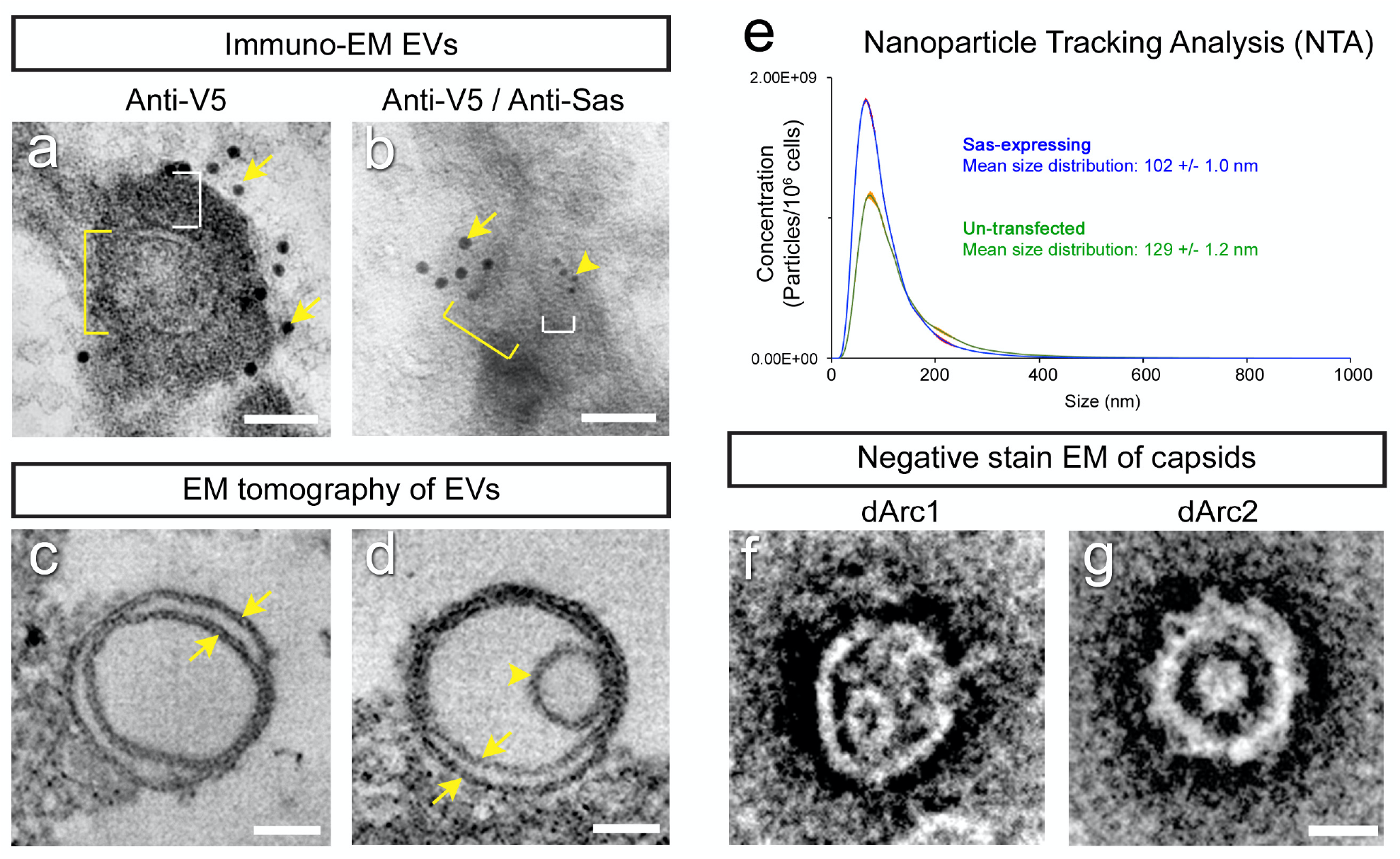
Analysis of EVs and capsids by electron microscopy and nanoparticle tracking analysis. **a, b**, Immuno-EM images of EVs from a purified EV prep from V5-Sas^FL^-expressing S2 cells. EV outline (membrane) diameters are indicated by yellow brackets. White brackets, separation between EV outline and a gold particle. **a**, immuno-EM with 10 nm anti-V5 gold particles (arrows). **b**, immuno-EM with both 10 nm anti-V5 (large gold, arrow) and 5 nm anti-Sas (small gold, arrowhead). **c**, EM tomogram of an empty double-membrane vesicle (arrows). Apparent EV sizes differ between immuno-EM and tomography, which use very different preparation methods. A low-mag view of a single slice from an EM tomogram of an EV preparation is shown in Supp. Fig. 2. **d**, EM tomogram of a double-membrane vesicle (arrows) with a capsid-sized denser object inside it (arrowhead). Video 1 shows a 3D reconstruction of this EV. Empty and filled single-membrane EVs were also observed (Supp. Fig. 2). Scale bars in **a-d**, 50 nm. **e**, Nanoparticle Tracking Analysis of purified EV preparations from untransfected (green curve) and Sas^FL^-expressing (blue curve) S2 cells. The mean size distribution is indicated. Standard error indicated by red color around curves. **f-g**, negative stain EMs of capsids from purified dArc1(**f**) and dArc2 (**g**) preparations from *E. coli*. Low-mag images of capsid preparations in Supp. Fig. 2. Scale bar in **f-g**, 20 nm. Source data files include an Excel file of raw data for the NTA analysis, the conversion of the numbers from numbers of EVs per sample to numbers of EVs per cell, based on cell counts, and plots of the data.

To analyze the numbers and sizes of EVs from Sas^FL^-expressing and control S2s, we examined purified EVs using Nanoparticle Tracking Analysis (NTA, System Biosciences, LLC). We observed that the distribution of EV diameters is shifted toward smaller values in the cells expressing Sas^FL^ (mean diameter=102 nm *vs*. 129 nm for control cells) (Fig. 2e). The mode (most frequently observed EV size) in the Sas^FL^ cells is about 70 nm, which is consistent with the diameters of many of the EVs we observed by EM tomography (Supp. Fig. 2). Expression of Sas^FL^ increased the number of EVs per cell in the exosome size range (30-160 nm in diameter) by 44%, and the number of EVs per cell of <100 nm in diameter by 72%, suggesting that the presence of high levels of Sas^FL^ increases the rate of EV production. This is consistent with a modest increase (∼60%) in the intensity of the Evi (EV marker) signal from Sas^FL^ expressing cells relative to control or Sas^short^ expressing cells that was observed in the Western blot experiment of Fig. 1c.

**Supp. Fig. 2.**
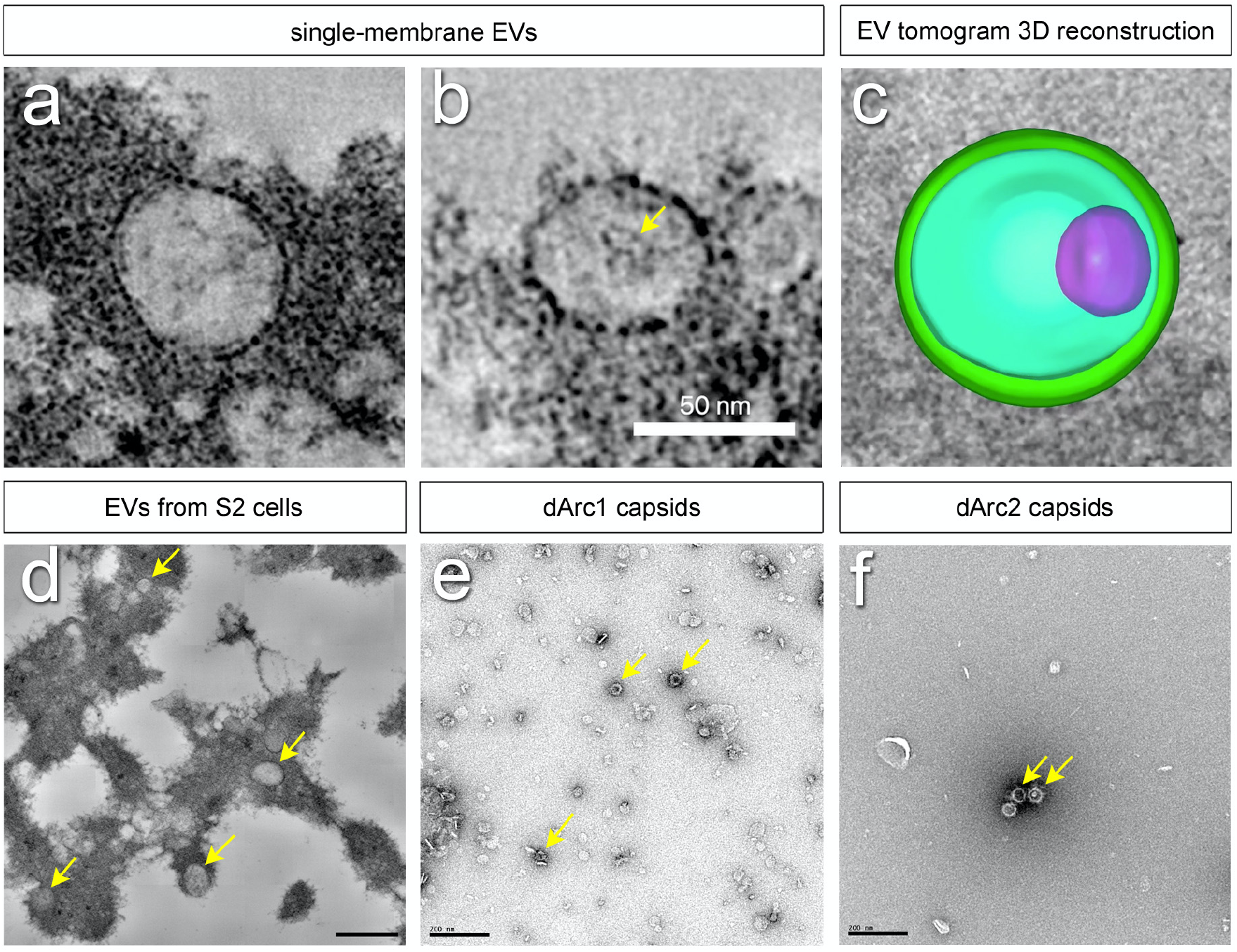
EM analysis of EVs from Sas^FL^-expressing S2 cells and dArc capsids. **a**, tomogram of an empty single-membrane EV. **b**, tomogram of a single-membrane EV enclosing a denser object (arrow). Scale bars in **a-b**, 50 nm. **c**, a still image from Video 1, which displays a reconstruction of the double-membrane EV from Fig. 2d. The denser object inside the EV is in magenta. **d**, a low-magnification image of a single slice from a tomogram, showing multiple EVs of various sizes (arrows). **e**, a low-magnification image of purified dArc1 capsids (arrows). **f**, a low-magnification image of purified dArc2 capsids (arrows). Scale bars in **d-f**, 200 nm.

### Sas^FL^ EVs target to cells expressing Ptp10D

Having shown that Sas^FL^ moves away from expressing neurons in the embryo and is an EV component, we then asked whether it can be incorporated into distant cells *in vivo*, presumably through endocytosis of EVs. We expressed V5-Sas^FL^ in 3^rd^ instar larval salivary glands (SGs) using an SG-specific GAL4 driver, *Sage-GAL4*, and visualized V5 staining in other tissues. We found that V5-Sas^FL^ made in SGs is present in imaginal discs, which are separated from SGs by larval hemolymph (Figs. 3a-b).

**Fig. 3.**
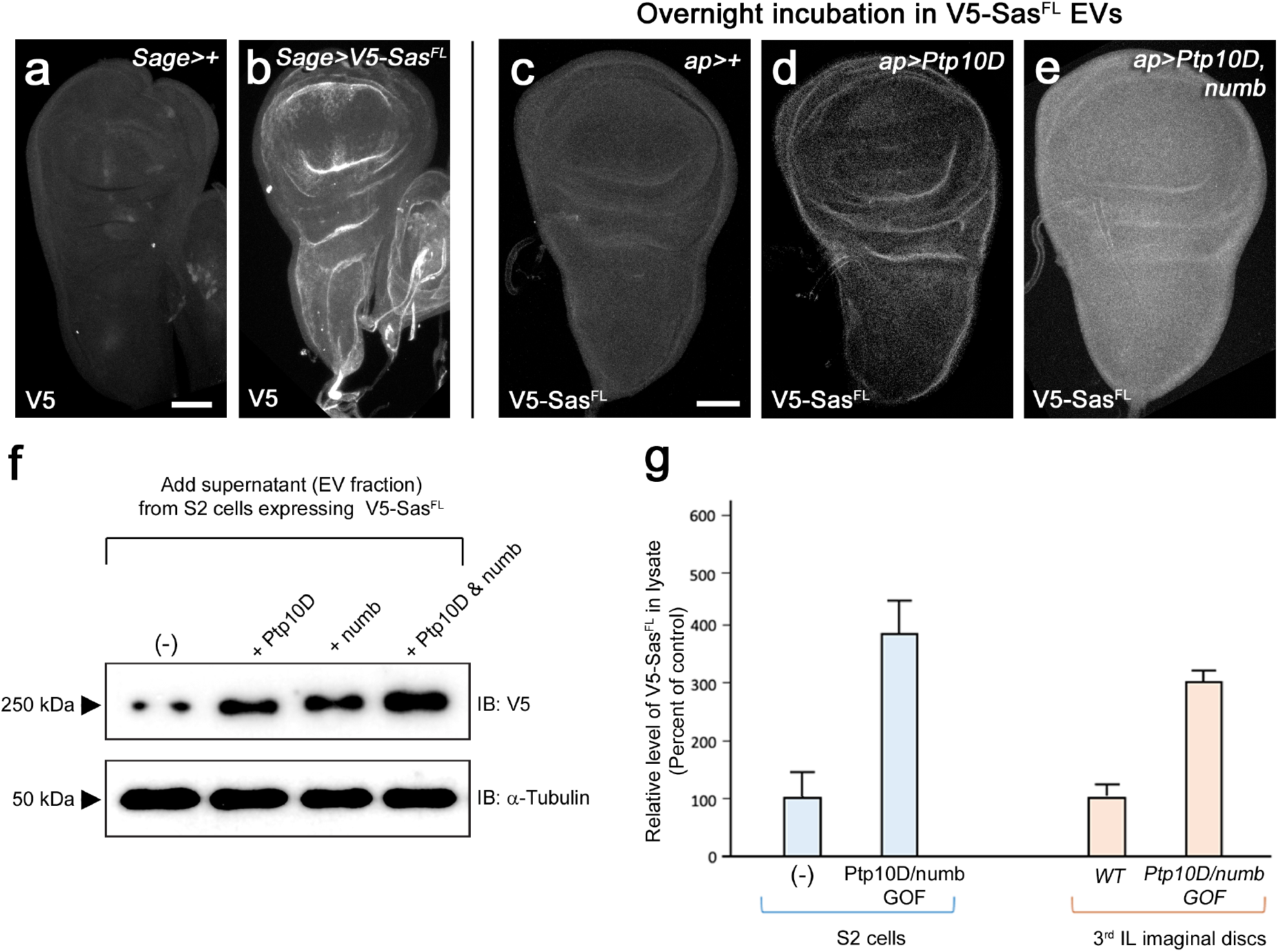
Transfer of Sas^FL^ to recipient cells. **a**, a third instar wing disc, with a portion of the haltere disc (right side), from a *Sage-GAL4/+* (SG-specific driver) larva, showing no V5 staining. **b**, a wing disc, with a portion of the haltere disc, from a *Sage>V5-Sas*^*FL*^ larva, showing bright V5 staining. Imaginal discs display no expression of GFP or mCherry reporters driven by *Sage-GAL4*. **c-e**, wing discs incubated with EVs from V5-Sas^FL^-expressing S2 cells and stained with anti-V5. For anti-Ptp10D and anti-Numb staining, see Supp. Fig. 3. **c**, ap-GAL4/+; **d**, ap>Ptp10D; **e**, ap>Ptp10D + Numb. **c**, low levels of anti-V5 staining are observed. **d**, higher levels are observed in disc folds, which also express Ptp10D (Supp. Fig. 3). **e**, bright anti-V5 staining is observed throughout the disc. This pattern matches anti-Numb staining (Supp. Fig. 3). Scale bars in **a** and **c**, 50 μm. **f**, transfer of Sas^FL^ from EVs into recipient S2 cells. Supernatants from S2 cells expressing V5-Sas^FL^ were incubated with cultures of untransfected S2 cells or cells expressing Ptp10D, Numb, or both, and cell lysates analyzed by Western blotting with anti-V5. Note that V5-Sas^FL^ levels were elevated relative to control cells by expression of either Ptp10D or Numb, and that levels were further increased by coexpression of Ptp10D and Numb coexpression. **g**, quantitation of results from panels **c-e** and **f**. Levels of transferred V5-Sas^FL^ were increased by ∼4-fold relative to untransfected controls by Ptp10D + Numb coexpression in S2 cells, and by ∼3-fold relative to ap-GAL4/+ control by Ptp10D + Numb coexpression in wing discs. Source data files include raw and labelled images for the Western blots shown in panel f, and an Excel file of the quantitation of the Western blot and disc immunofluorescence signals used to generate panel g.

To examine mechanisms involved in specific targeting of Sas EVs, we added supernatants (EV fraction) from V5-Sas^FL^-expressing S2 cells to S2 cell cultures and analyzed recipient cell lysates by Western blotting. We observed that expression of the Sas receptor Ptp10D in recipient cells increased V5-Sas^FL^ levels in these cells, as did expression of Numb, a regulator of endocytosis that binds to the Sas ICD(Chien *et al*., 1998). Expression of both Ptp10D and Numb produced a synergistic effect, increasing V5-Sas^FL^ by ∼4-fold relative to untransfected recipient cells (Figs. 3f-g). We speculate that binding of Numb to the Sas ICD increases Sas uptake and/or protects endocytosed Sas from degradation.

We then developed an assay to examine the effects of Ptp10D and Numb on Sas targeting in larval cells by incubating dissected 3^rd^ instar wing imaginal discs with V5-Sas^FL^ supernatants. We expressed Ptp10D, or both Ptp10D and Numb, in wing discs using the *Ap-GAL4* driver. The control wing discs displayed weak V5 staining after incubation with V5-Sas^FL^ EVs. Staining was increased by Ptp10D expression, and further elevated (∼3-fold increase relative to *Ap-GAL4* control) by expression of both Ptp10D and Numb (Figs. 3c-e, g; Supp. Fig. 3).

**Supp. Fig. 3.**
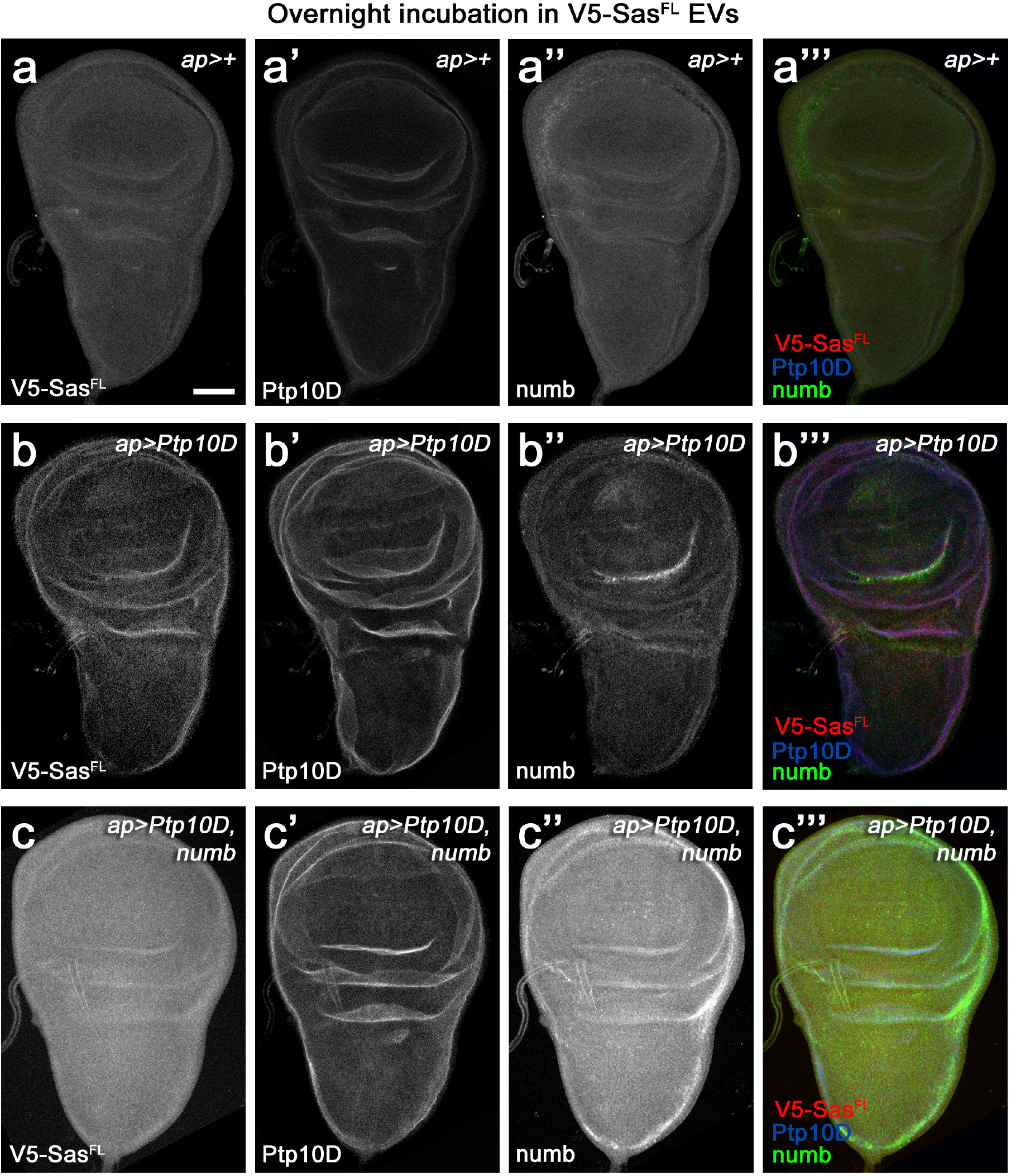
Visualization of all three channels of the triple-stained wing discs shown in Fig. 3. **a-c** show the anti-V5 channel, **a’-c’** the anti-Ptp10D channel, **a’’-c’’** the anti-Numb channel, and **a’’’-c’’’** all three channels. When Numb is overexpressed, it labels the entire disc (**c’’**). Ptp10D localizes to cells in disc folds when overexpressed (**b’** and **c’**). These folds label brightly with anti-V5 in **b**. Ap-GAL4 expresses GFP reporters in the dorsal ¾ of the disc, but not in the ventral region. However, in discs incubated overnight with supernatant, we do not observe clear borders in the Numb and Ptp10D expression patterns.

### Sas binds to dArc1 and mammalian Arc via a conserved tyrosine motif

We then examined whether Sas interacts with specific EV cargoes. To do this, we made EV preparations from S2 cells expressing V5-Sas^FL^ and from untransfected control cells, lysed them with nonionic detergent, incubated the lysates with anti-V5-coupled magnetic beads, and analyzed bead-bound proteins by mass spectrometry (Fig. 4a, Supp. Table 1). We ranked the identified proteins by their degree of enrichment in the V5-Sas^FL^ samples relative to controls. Proteins that are present in the V5-Sas^FL^ samples should include EV cargoes that bind to Sas^FL^ and are therefore present in V5 IPs. Proteins in control samples would be those that nonspecifically bind to V5 beads. We observed that the most highly enriched protein (after Sas itself) is dArc1 (22-fold) (Fig. 4b). dArc2 is #7 on the list (6-fold). *dArc1* mRNA, presumably encapsulated within dArc1 capsids, is known to be a prominent mRNA component of EVs from *Drosophila* cultured cells(Ashley *et al*., 2018; Lefebvre et al., 2016). We then went on to show that dArc1 binds directly to the Sas ICD (see below).

**Fig. 4.**
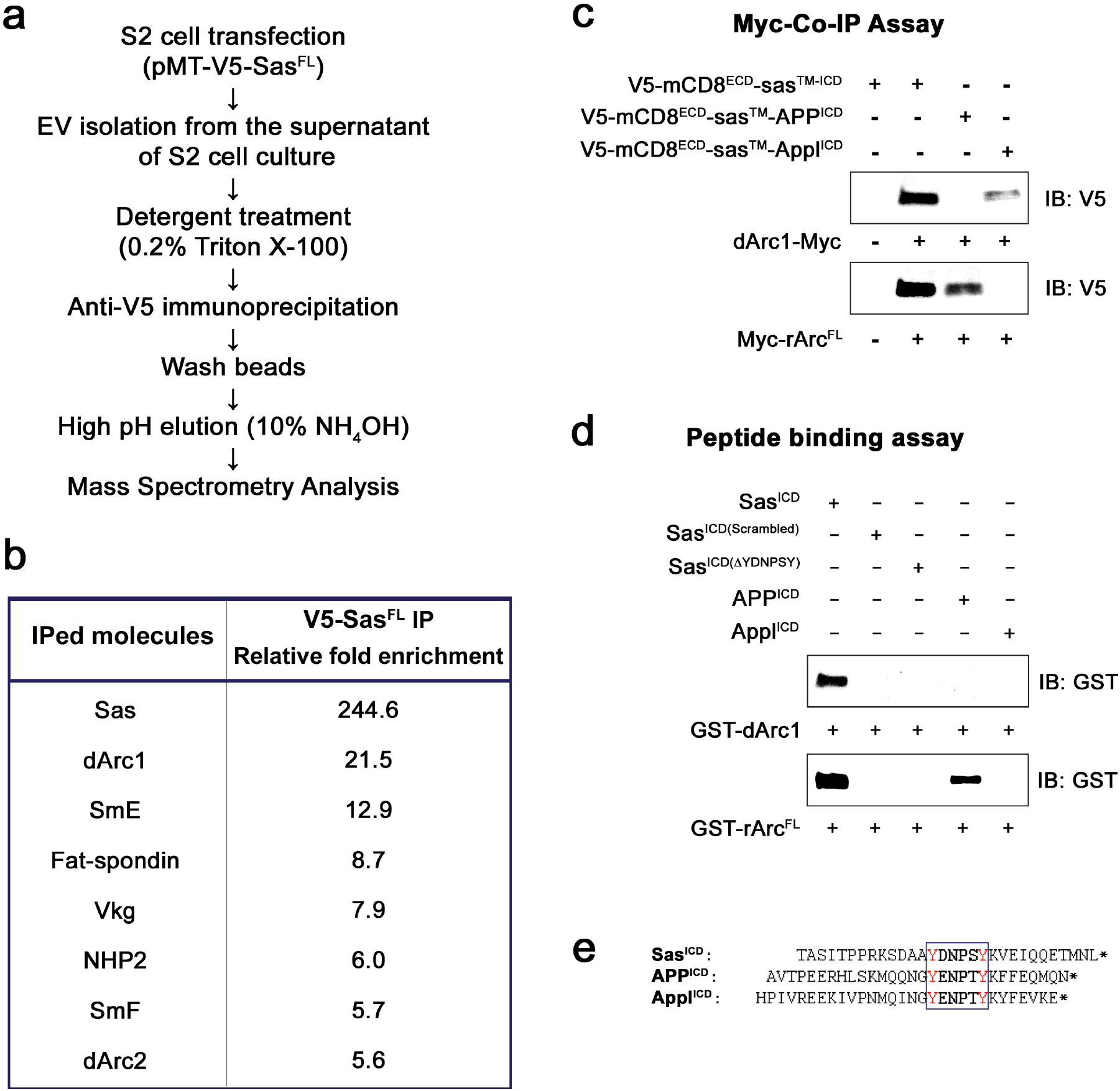
Interactions of Sas, Appl, and APP with Arcs. **a**, protocol for mass spectrometry analysis. Purified EVs from control S2 cells or S2 cells expressing V5-Sas^FL^ were lysed and IP’d with anti-V5, followed by protease digestion and mass spectrometry analysis. **b**, mass spectrometry results. The 7 proteins present at the highest levels in IPs from V5-Sas^FL^ EVs relative to IPs from control EVs (> 6-fold ratio) are listed. Sas itself was the most highly enriched protein, as expected. dArc1 and dArc2 were enriched by 22-fold and 6-fold, respectively. **c**, co-IP/Western blot analysis of association between Sas and Arc fusion proteins in transfected S2 cells. S2 cells were transfected with the V5-mCD8^ECD^-Sas^TM-ICD^ fusion protein construct, or with equivalent constructs in which the Sas ICD was replaced by the Appl or APP ICD, with or without Myc-tagged dArc1 or mammalian (rat) Arc (rArc^FL^) constructs. Lysates were IP’d with anti-Myc and blotted with anti-V5. Anti-V5 bands of the correct size were observed when dArc1 was expressed with Sas or Appl ICD constructs, and when rArc^FL^ was expressed together with Sas or APP ICD constructs. **e**, direct binding of purified GST-dArc1 and GST-rArc^FL^ fusion proteins to Sas, APP, and Appl ICD peptides. Biotinylated peptides were bound to streptavidin magnetic beads, which were incubated with GST-Arc proteins, followed by Western blotting of bead-bound proteins with anti-GST. dArc1 bound to the wild-type, but not to scrambled or YDNPSY deletion mutant Sas ICD peptides, while rArc^FL^ bound to wild-type Sas and APP ICD peptides. **f**, sequences of the complete Sas, APP, and Appl ICDs, corresponding to biotinylated peptide sequences. The conserved tyrosine motif is boxed, with tyrosines in red. *, stop codons.

Other proteins within the top 7 included small ribonucleoproteins (SmE and SmF), a ribosomal protein (NHP2), and a collagen (Vkg). Proteins in these categories were found to be major EV components in a proteomic analysis of S2 and Kc167 EVs(Koppen et al., 2011). We think it likely that some or all of these proteins are abundant contaminants that do not actually interact with Sas but happened to be present at higher levels in the IP from Sas-expressing cells *vs*. the IP from control cells. We did not further examine any of these proteins.

EVs from media of short-term cultures of mouse cortical neurons were shown to contain denser objects whose size (∼30 nm in diameter) was consistent with mammalian Arc capsids, and which were associated with anti-Arc gold particles(Pastuzyn *et al*., 2018). For dArc1, capsid-like structures that bound to anti-dArc1 gold particles were detected in lysed preparations of EVs from S2 cells(Ashley *et al*., 2018). We examined the EVs from Sas^FL^-expressing S2 cells EVs by EM tomography, and were able to visualize denser objects within many of them (Fig. 2d, Supp. Fig. 2b). These were ∼40 nm in diameter, consistent with the known dimensions of the dArc1 capsid (37 nm)(Erlendsson *et al*., 2020; Hallin et al., 2021). A video of a 3D reconstruction of the EM tomogram of the EV in Fig. 2d is included (Video 1), and Supp. Fig.2c shows a still image from this video.

Since dArc1 was enriched in Sas^FL^ preparations purified from EV lysates with anti-V5, we then investigated whether it binds to the Sas ICD (which would be in the EV interior) by co-IP in S2 cells. We coexpressed Myc epitope-tagged dArc1 with a fusion protein in which the V5-tagged ECD of mouse CD8 was attached to the TM domain and the 37 aa ICD of Sas. We then IP’d cell lysates with anti-Myc, and detected V5-mCD8^ECD^-Sas^TM-ICD^ by Western blotting. We observed that purified dArc1 co-IP’d with the Sas ICD fusion protein (Fig. 4d). We performed the same experiment for dArc2, but did not observe a consistent co-IP signal.

The Sas ICD sequence contains the sequence motif YDNPSY, which is a PTB-binding motif (NPXY) that overlaps by two amino acids with an SH2-binding motif (YXXP) that is also a potential Abl tyrosine kinase substrate sequence(Colicelli, 2010) (Fig. 4e). The NPXY motif is the target for binding of the Numb PTB(Li et al., 1998). This suggests that an SH2 protein and a PTB protein might compete for binding to this sequence, if the first tyrosine was phosphorylated to create an SH2 docking site. The PTB domain of Numb does not require tyrosine phosphorylation to bind to its NPXY target. Interestingly, in an earlier mass spectrometric analysis, we found that the Shc protein, which contains a phosphotyrosine-binding SH2 domain, was associated with Sas purified from S2 cells treated with pervanadate to induce high-level tyrosine phosphorylation.

We searched for other *Drosophila* CSPs containing a sequence with similar properties in their ICDs, and found only one, Appl, which has the sequence YENPTY but is otherwise unrelated to the Sas ICD. Human APP, the mammalian ortholog of Appl, contains the same sequence in its short ICD (Fig. 4e), as do the two APP paralogs, APLP1 and APLP2. We then replaced the Sas ICD in the V5-mCD8^ECD^-Sas^TM-ICD^ construct with the Appl and APP ICDs, and found that the Appl ICD protein co-IP’d with dArc1 (Fig. 4d), implicating the Y(D/E)NP(S/T)Y sequence in binding to dArc1. Interestingly, this sequence contains the consensus motif for binding of mammalian Arc to TARPγ2, CaMKII, and NMDA receptor peptides, which is X-P-X- (Y/F/H)(Nielsen et al., 2019; Zhang et al., 2015). Arc binds to the NMDA receptor as a monomer(Nielsen *et al*., 2019). The TARPγ2 Arc-binding peptide is RIPSYR, which is similar to the sequences in Sas (PSYK) and APP (PTYK). Accordingly, we expressed Myc-tagged mammalian Arc (rArc^FL^) in S2 cells and examined whether it could co-IP with the V5-mCD8-ICD fusion proteins. We observed that Arc was able to co-IP with the Sas and APP ICDs (Fig. 4d). This is interesting, because mammalian and *Drosophila* Arc are not orthologs, and are apparently derived from independent Ty3/gypsy retrotransposon lineages(Ashley *et al*., 2018; Hantak *et al*., 2021; Pastuzyn *et al*., 2018). The fact that both proteins mediate intercellular communication suggests that they may be products of convergent evolution. Fly and mammalian Arc appear to have evolved preferences for binding to similar peptide sequences.

The co-IP data indicate that the Sas ICD associates with dArc1 and Arc, but does not show that the two proteins directly interact. To evaluate this, we made the complete Sas, APP, and Appl ICDs (Fig. 4e), as well as a scrambled version of the Sas ICD and a deletion mutant of the Sas ICD that lacks the YDNPSY sequence, as biotinylated peptides, and bound these to streptavidin-coupled magnetic beads. To make purified Arc proteins for binding, we expressed dArc1, dArc2, and mammalian Arc as GST fusion proteins in *E. coli*. To evaluate the properties of these proteins, we cleaved off the GST after purification to facilitate capsid formation(Nielsen *et al*., 2019) and visualized the preparations by negative-stain EM. The dArc1 and dArc2 preparations contained ∼40 nm diameter capsids that appeared similar to those observed in previous studies(Ashley *et al*., 2018; Erlendsson *et al*., 2020; Pastuzyn *et al*., 2018) (Figs. 2f-g).

We then mixed the beads with purified GST-dArc1, GST-rArc^FL^, and GST-dArc2 proteins and examined whether we could observe specific binding. As a positive control, we made purified Numb PTB domain, and showed that it bound as expected to the Sas, APP, and Appl peptides, which all contain the NPXY PTB-binding motif, but not to the scrambled Sas peptide or the YDNPSY deletion mutant. In the peptide binding assay, we observed that dArc1 directly bound to the wild-type (wt) Sas ICD sequence, but not to the other peptides. Mammalian Arc also bound to the wt Sas ICD, as well as to the APP ICD (Fig. 4e). GST-dArc2 did not bind specifically to any peptides.

These results implicate the Y(D/E)NP(S/T)Y sequence as a determinant of binding to Arcs (Fig. 4e). The data suggest that APP might be a CSP that has a relationship to Arc which is similar to that of Sas to dArc1. This will be of interest to explore in future studies, especially since Arc has been implicated in AD pathogenesis(Bi *et al*., 2018; Landgren *et al*., 2012; Wu *et al*., 2011). The first Y in the YENPTY motif in APP has been reported to be a substrate for the Abl tyrosine kinase(Zambrano et al., 2001). If YENP was phosphorylated, it would become a docking site for a class of SH2 domain proteins, and binding of this protein(s) could occlude Arc binding to the adjacent PTYK sequence. The Abl inhibitor imatinib (Gleevec), which would be expected to block phosphorylation of this site, inhibits formation of β-amyloid peptide (Aβ)(Netzer et al., 2003), and binding of Arc to APP could be relevant to this effect.

### Sas facilitates intercellular transfer of dArc1 and its mRNA *in vivo*

Sas is not required for loading of dArc1 capsids into EVs, since *dArc1* mRNA is a normal component of EVs from cell lines that do not express Sas. If it behaves like mammalian Arc in its interactions with peptides(Nielsen *et al*., 2019), dArc1 might bind to Sas as a monomer. Perhaps Sas recruits dArc1 monomers (possibly bound to mRNA *via* their Zn^2+^ fingers) to nascent EVs during their biogenesis, and they then assemble into capsids. Binding of Sas to dArc1 may help to increase the probability that Sas-bearing EVs contain dArc1 capsids. The function of Sas would then be to deliver the EVs and their dArc1 capsid cargo to specific recipient cells.

Having shown that Sas^FL^ can move within larvae and that it binds to dArc1, which is a known component of EVs that mediates intercellular communication, we then examined whether it can cause dArc1 to move from source cells into recipient cells *in vivo*. To establish an assay system for dArc1 capsid movement, we first expressed V5-Sas^FL^ in late stage 16 embryonic SGs together with RFP, and observed that V5 signal moved to the gut and tracheae, while RFP was retained in the SGs as expected (Figs. 5a-b).

**Fig. 5.**
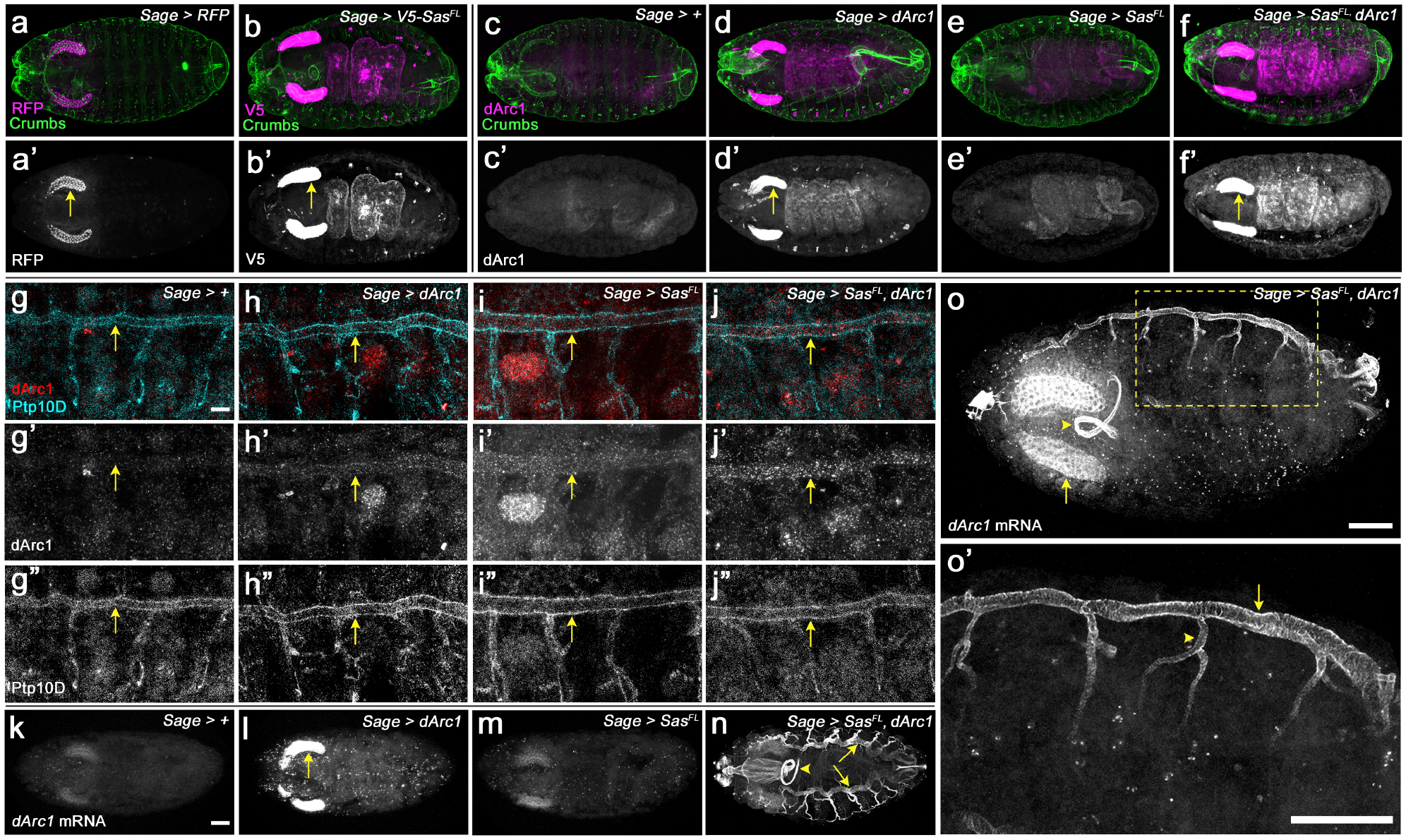
Sas facilitates transfer of dArc1 capsids bearing dArc1 mRNA into distant cells *in vivo*. **a-b**, localization of RFP (**a**) and V5-Sas^FL^ (**b**) driven by Sage-GAL4. Whole-mount late stage 16 embryos (top-down view, anterior to the left) were double-stained with anti-RFP (**a**) or anti-V5 (**b**) (magenta) plus anti-Crumbs (apical marker, expressed in epithelia, including tracheae; green). **a’** shows the RFP channel alone, and **b’** shows the V5 channel. Arrows, SGs. Note that V5-Sas^FL^ is observed in the gut and peripheral dots, while RFP is retained in the SGs. **c-f**, localization of dArc1 protein in whole-mount late stage 16 embryos. **c**, control (*Sage-GAL4/+*); **d**, *Sage>dArc1*; **e**, *Sage>Sas*^*FL*^; **f**, *Sage>Sas*^*FL*^ *+ dArc1*. **c-f** show double-staining with anti-dArc1 (magenta) and anti-Crumbs (green). **c’-f’** show the dArc1 channel alone. Arrows, SGs. Bright dArc1 SG staining is observed when dArc1 is expressed. When Sas^FL^ and dArc1 are both expressed, bright dArc1 staining of the gut and peripheral dots is observed. Weaker gut staining is observed when dArc1 is expressed alone. **g-j**, localization of dArc1 protein and Ptp10D in high-magnification views of body walls from fillets of late stage 16 embryos (anterior to the left, dorsal up). **g**, control (*Sage-GAL4/+*); **h**, *Sage>dArc1*; **i**, *Sage>Sas*^*FL*^; **j**, *Sage>Sas*^*FL*^ *+ dArc1*. **g-j** show double-staining with anti-dArc1 (red) and anti-Ptp10D (blue). **g’-j’** show the dArc1 channel alone. **g’’-j’’** show the Ptp10D channel alone. Arrows, dorsal tracheal trunk. There are numerous bright dArc1 puncta in the tracheal trunk when Sas^FL^ and dArc1 are expressed together. Fewer and weaker puncta are observed when Sas^FL^ or dArc1 are expressed alone, and no puncta are seen in *Sage-GAL4/+* controls. **k-n**, *dArc1* mRNA from the endogenous gene, detected by FISH with a 3’ UTR probe. **k**, control (*Sage-GAL4/+*); **l**, *Sage>dArc1*; **m**, *Sage>Sas*^*FL*^; **n**, *Sage>Sas*^*FL*^ *+ dArc1*. There is weak expression of dArc1 mRNA in the SGs in controls. When dArc1 (from an ORF construct) is expressed alone, bright SG staining is observed, indicating that exogenous dArc1 increases expression of endogenous *dArc1* mRNA. There are also scattered *dArc1* mRNA puncta elsewhere in the embryo. When Sas^FL^ and dArc1 are expressed together, bright *dArc1* mRNA FISH staining of the entire tracheal system is observed (arrows indicate dorsal tracheal trunks), as well as the foregut (arrowhead) and esophagus. **o, o’**, high-magnification views of *dArc1* mRNA in the tracheae in an obliquely mounted (anterior to the left, dorsal up) embryo expressing Sas^FL^ and dArc1. **o’** is a higher-magnification inset (yellow dotted outline) from **o**. Arrow in **o**, SG; arrowhead, foregut loop. Arrow in **o’**, dorsal tracheal trunk; arrowhead, transverse connective. Scale bar in **k** (applies to **a-f** and **k-n**), 50 μm; scale bar in **g** (applies to **g-j**), 10 μm; scale bar in **o**, 50 μm; scale bar in **o’**, 50 μm.

To examine dArc1 transport, we needed to express untagged dArc1 and visualize it with antibody against dArc1(Ashley *et al*., 2018), because we were unsuccessful in detecting movement of tagged versions of dArc1. dArc1 is made at very low levels in embryos. In late stage 16 control embryos (*Sage-GAL4/+*), we observed faint ubiquitous staining, with higher levels in the gut. The same pattern was observed when Sas^FL^ alone was expressed in SGs, although gut staining was slightly increased (Figs. 5c’, 5e’). We then expressed dArc1 from a UAS construct that contained only the dArc1 open reading frame (ORF), flanked by heterologous 5’ and 3’ UTR sequences. The short 3’ UTR was derived from SV40. When we expressed dArc1 alone in SGs, we observed bright anti-dArc1 staining in the SGs and increased staining relative to controls in the gut and in dots in the body wall (Fig. 5d’). Expression of both Sas^FL^ and dArc1 produced a larger increase in dArc1 staining in the gut and peripheral dots (Fig. 5f’).

To localize dArc1 staining in the body wall and compare it to Ptp10D staining, we examined dissected “fillets” at high magnification. For reference, Supp. Figs. 4a-d show the evolution of Ptp10D expression from stage 14 to late stage 16. VNC expression continuously increases during this time period, while tracheal expression begins in stage 14, decreases in stage 15, and re-emerges at stage 16, at which time Ptp10D is expressed in the main tracheal trunk and major tracheal branches. Fig. 4j’ shows that, in late stage 16 embryos expressing both Sas^FL^ and dArc1 in SGs, there were many bright puncta stained with anti-dArc1 in the dorsal tracheal trunk, which expresses Ptp10D. These puncta appeared similar to those previously observed at larval NMJs(Ashley *et al*., 2018). They were not detectable in control embryos (*Sage-GAL4/+*).

There are lower numbers of fainter dArc1 puncta in tracheal trunks of the two other genotypes (*Sage>dArc1* and *Sage>Sas*^*FL*^)(Figs. 5h’, i’). Endogenous Sas is expressed at low levels in SGs, and endogenous *dArc1* mRNA is also present in SGs (Fig. 5k), although dArc1 protein is not detectable. Endogenous Sas^FL^ may be able to transport some of the overexpressed dArc1, and overexpressed Sas^FL^ might transport some endogenous dArc1, giving rise to the observed puncta. It is also interesting that dArc1 (and *dArc1* mRNA; see below) is observed in tracheal cells, but not in VNC neurons, which also express Ptp10D at high levels. There is a glial sheath around the VNC at late stage 16, and this might block access of EVs to Ptp10D-expressing neurons. Alternatively, perhaps there are cofactors required for EV binding and/or internalization that are not expressed in neurons.

More dramatic effects of Sas^FL^ on dArc1 capsid movement were observed when endogenous *dArc1* mRNA was examined by fluorescence *in situ* hybridization (FISH). To detect mRNA, we used the 700 nt antisense 3’ UTR probe employed in the (Ashley *et al*., 2018) paper to visualize *dArc1* mRNA puncta at the NMJ. Note that this probe does not recognize overexpressed *dArc1* mRNA made from the UAS construct, because that contains only the dArc1 ORF and no *dArc1* 3’ UTR sequences. In late stage 16 control embryos (*Sage-GAL4/+*), we observed faint FISH signals in the SGs and a few puncta elsewhere in the embryo (Fig. 5k). A similar pattern was seen in *Sage>Sas*^*FL*^ embryos (Fig. 5m). However, when dArc1 was expressed from the UAS-dArc1 ORF construct, we observed bright FISH signals in SGs with the 3’ UTR probe (Fig. 5l). There were also scattered puncta in other parts of the embryos. This shows that exogenous dArc1 induces expression of endogenous *dArc1* mRNA (or stabilizes the mRNA). No signal was observed when a sense *dArc1* probe was used for FISH (Supp. Figs. 4e-h). Finally, when Sas^FL^ and dArc1 were expressed together, we observed a completely different pattern, in which the entire tracheal system is lit up by the FISH signal (Fig. 5m). The foregut and esophagus also stain brightly.

Figs. 5o and 5o’ show the tracheae and SGs at higher magnification, in side views of an embryo expressing both Sas^FL^ and dArc1 in SGs. The dorsal tracheal trunk (arrow) and the transverse connective (arrowhead) both display bright *dArc1* FISH signals. Note that, because this is a confocal image (optical section), the cells at the edges of the tracheal trunk are bright, while the hollow lumen is dark. The brightness of the tracheal FISH signal suggests that it represents not only *dArc1* mRNA transferred from capsids, but *dArc1* mRNA synthesized in these cells in response to dArc1 protein made from the transported capsid mRNA. If this is correct, it would represent an amplification mechanism in which translated *dArc1* mRNA from EVs can induce expression of much more *dArc1* mRNA in the recipient cells. Finally, we examined whether the Sas ICD is required for *dArc1* mRNA transport by expressing dArc1 together with a protein (Sas^ECD-TM^-GFP) in which the Sas ICD was replaced by GFP. This protein is present in EVs when expressed in S2 cells, but it does not produce any *dArc1* FISH signal outside of the SGs (Supp. Fig. 4i), indicating that it cannot facilitate transport of dArc1 capsids to tracheal cells.

**Supp. Fig. 4.**
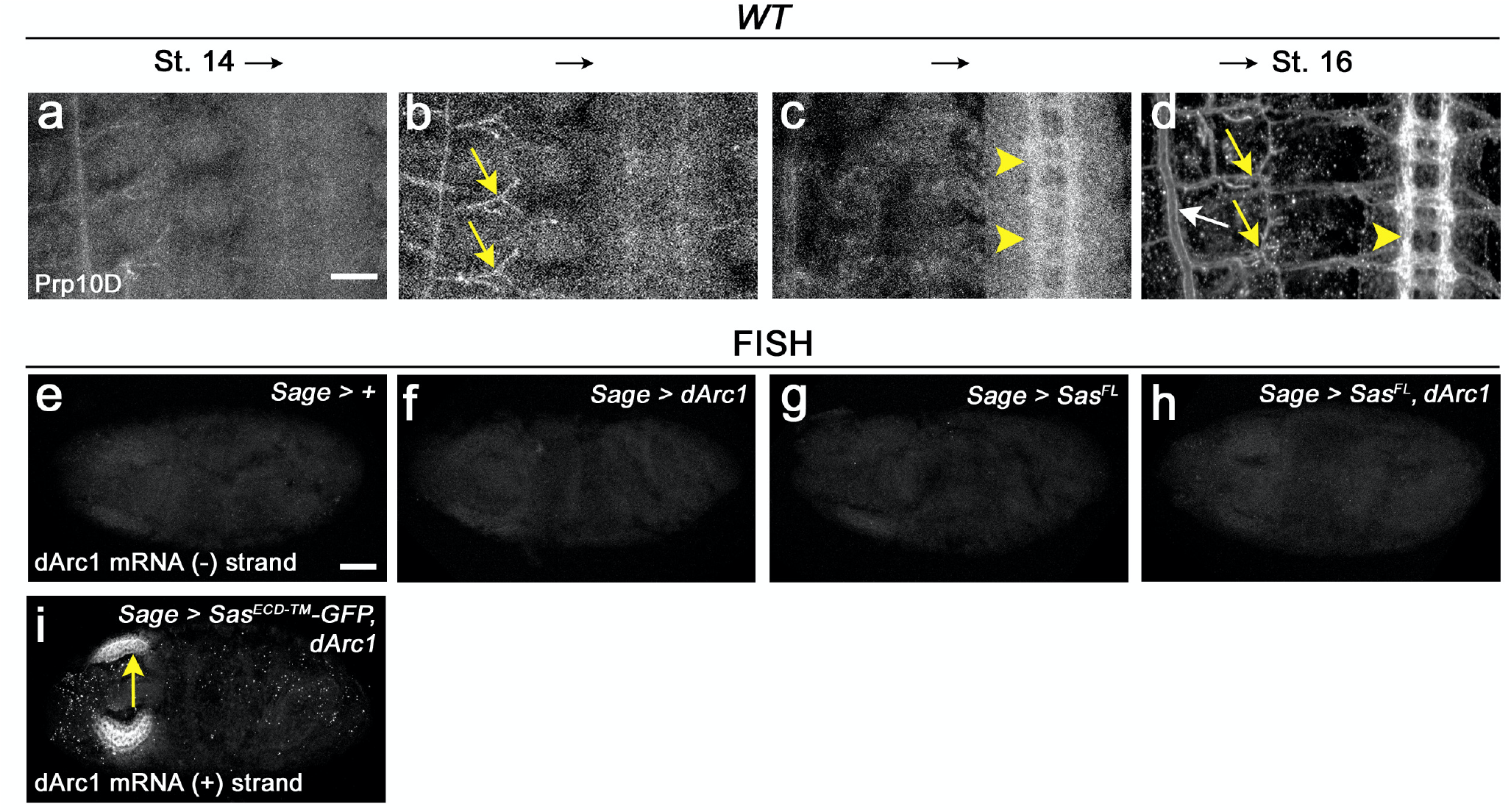
Ptp10D expression, *dArc1* sense control FISH data, and *dArc1* antisense FISH data from embryos expressing Sas^ECD-TM^-GFP and dArc1. **a-d**, expression of Ptp10D in the tracheae (arrows) and VNC axons (arrowheads) of embryos of advancing age, from stage 14 (**a**), through stage 15 and early 16 (**b** and **c**), to late stage 16 (**d**). In **d**, the white arrow indicates the dorsal tracheal trunk and the yellow arrow indicates the transverse connective. **e-h**, FISH results with the *dArc1* sense probe control for the four genotypes. No specific staining is observed. **i**, FISH analysis of *dArc1* mRNA expression in embryos expressing Sas^ECD-TM^-GFP and dArc1. The arrow indicates an SG, which stains brightly. There are only scattered puncta elsewhere in the embryo.

## Conclusions

Our results on movement of Sas EVs containing dArc1 capsids are summarized in the diagram of Fig. 6. These findings contribute to the understanding of intercellular communication mechanisms by showing that Sas is an EV targeting ligand that directs internalization of EVs into cells expressing the Sas receptor Ptp10D. dArc1 is related to retrotransposon Gag proteins, and it forms a capsid that contains *dArc1* mRNA and is loaded into EVs(Ashley *et al*., 2018). Sas facilitates transfer of dArc1 capsids into Ptp10D-expressing recipient cells *in vivo*. The Sas ICD binds directly to dArc1. Mammalian Arc also forms capsids that are transported *via* EVs(Pastuzyn *et al*., 2018), and it binds to the Sas and APP ICDs, which share a tyrosine motif. The connection between Arc and APP will be of interest to explore in future studies, because Arc has been linked to β-amyloid accumulation and AD pathogenesis(Bi *et al*., 2018; Landgren *et al*., 2012; Wu *et al*., 2011). Also, full-length APP and some of its proteolytic products are localized to EVs, and EVs from N2a cells bearing tagged APP are internalized into cultured neurons, but not into glia(Laulagnier *et al*., 2018). It will be interesting to determine if APP EVs contain Arc capsids, and if the presence of APP on Arc-containing EVs causes Arc to be preferentially delivered to a specific population of neurons.

**Fig. 6.**
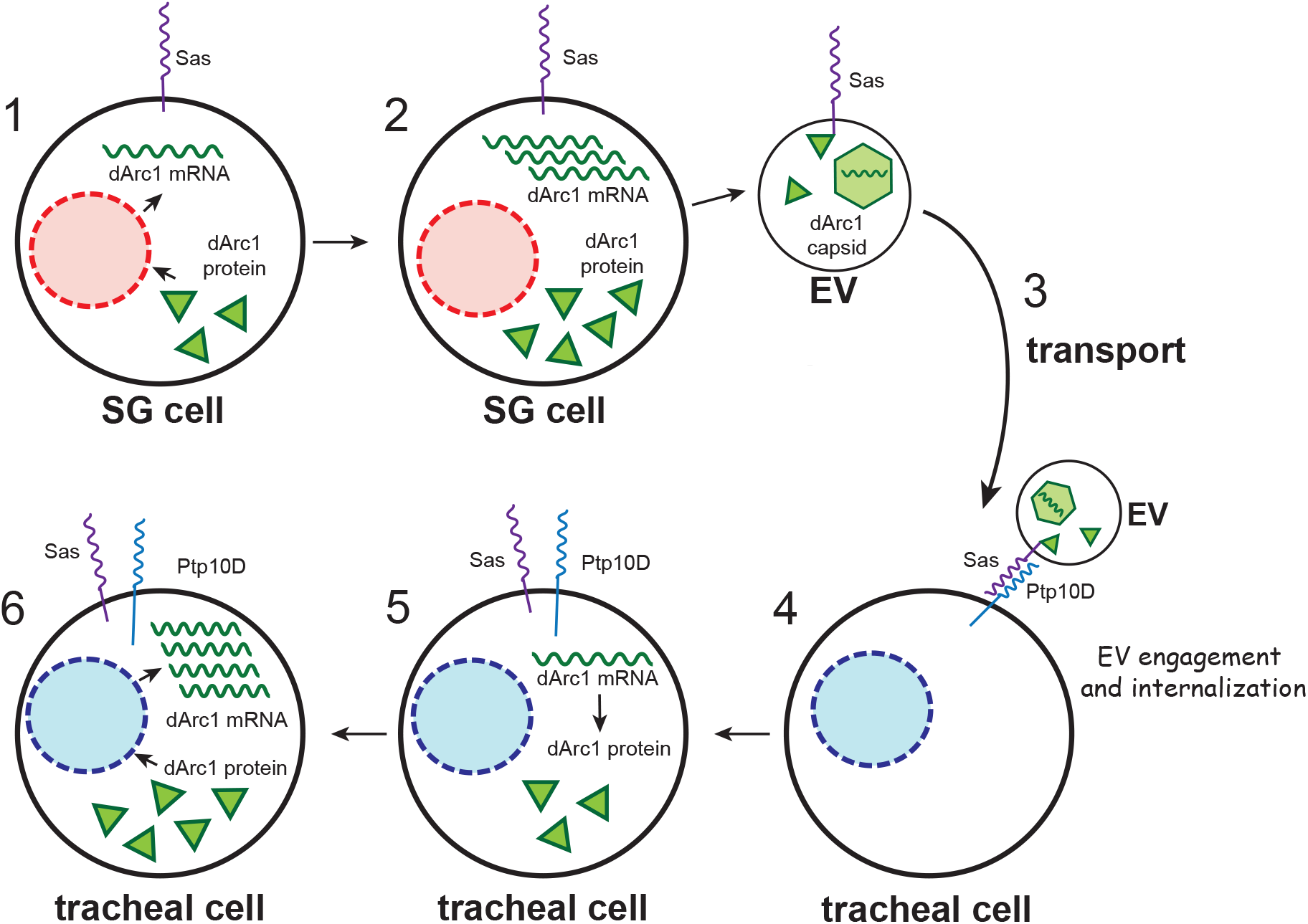
Schematic diagram of the processes involved in movement of EVs bearing Sas and dArc1 capsids from salivary glands to tracheal cells. Steps 1 and 2, expression of the dArc1 ORF induces accumulation of *dArc1* mRNA in SGs. EVs with Sas^FL^ on their surfaces bearing dArc1 capsids diffuse or are transported through the hemolymph (Step 3) and bind to Ptp10D-expressing tracheal cells (Step 4). The EVs internalize into the tracheal cells and release *dArc1* mRNA (Step 5), and dArc1 protein induces high-level expression of more *dArc1* mRNA.

## Methods

### Fly stocks and genetics

The following stocks were used: *yw* for wild-type control, *ap-GAL4* (Bloomington 50156), *UAS-mCD8::GFP* (Bloomington 5130), *UAS-myr::mRFP* (Bloomington 7118), *UAS-mCherry*.*NLS* (Bloomington 38424), *sas15* (null mutant)(Bloomington 2098), *Sage-GAL4* (a gift from Deborah J. Andrew), *Ptp10DEP1172* (Bloomington 11332), *UAS-dArc1* (Bloomington 37532), *UAS-Numb* (a gift from Yuh Nung Jan), *UAS-SasFL* and *UAS-V5-SasFL* (Lee et al., 2013), *Arc1esm18* (Bloomington 37530). Crosses and embryo collections were performed at room temperature. For overexpression experiments, embryos were shifted to 29°C for at least 120 min prior to fixation and staining and 3^rd^ instar larvae were shifted to 29°C for overnight for further analysis. For the EV targeting experiments (Figs. 3c-e), imaginal discs from 3^rd^ instar larvae were harvested at room temperature and incubated in 200 μl of S2 supernatant overnight at 29°C before fixation and staining. There are 10,000-50,000 cells in a 3^rd^ instar imaginal disc. Given the results from the NTA analysis, we can conclude that ∼140,000 EVs are present in 200 μl of supernatant from V5-Sas^FL^-expressing S2s cells. We used 5 wing discs per incubation, so the ratio of EVs to cells is ∼0.5 to ∼2. The relative V5 signal intensities on the imaginal discs were measured by densitometry analysis using ImageJ software.

### Immunohistochemistry

Embryos and larval tissues were stained with standard immunohistochemical procedures. The following antibodies were used: rabbit anti-V5 (1:1,000, Invitrogen); mouse anti-GFP (1:1,000, Invitrogen); rabbit-anti-Sas^FL^ (1:2,000, gift of D. Cavener); rat-anti-Sas^short^ (1:50, GenScript USA Inc.); mAb Cq4 against crumbs (1:100, DSHB); guinea pig-anti-Numb (1:1,000, gift from J. Skeath); rabbit-anti-dArc1 (1:100, gift from T. Thomson); mAb 8B2 against Ptp10D (1:5, DSHB); mAb MR1A against Prospero (1:40, DSHB); rat-anti-Repo (1/2,000, gift from S. Banerjee); rabbit anti-Evi (Wntless, 1:5000, gift from K. Basler); FITC-conjugated phalloidin (1:1,000, Thermo Fisher Scientific); AlexaFluor 488 anti-mouse, AlexaFluor 488 anti-rat, AlexaFluor 568 anti-rabbit, AlexaFluor 568 anti-rat and AlexaFluor 647 anti-mouse (1:1,000, Invitrogen). Rat anti-Sas^short^ antibody was generated against a synthetic peptide, HSSIPANGANNLQP, flanking the EVT region (intron is between the N and G residues) and the KLH-conjugated antibody was purified by protein G column (GenScript USA Inc.). Samples were mounted in VECTASHIELD (Vector Laboratories) and analyzed on a Zeiss LSM 880.

### Cell culture and preparation of EVs and cell lysates

EVs and cell lysates were prepared from S2 cells that were cultured for four days at 22°C in Schneider’s medium (Gibco) supplemented with 10% exosome-free FBS (#EXO-FBSHI-50A-1, SBI) to avoid contamination from Bovine serum exosomes. DNA constructs were transiently transfected into S2 cells using Effectene (Qiagen). EVs for Western blot analysis (Fig. 1c) and electron microscopy (Figs. 2 and Supp. Fig.) were collected using Total Exosome Isolation reagent (#4478359, Invitrogen) from the supernatants of S2 cultures. This kit has been found to produce exosomes of equivalent quality from mammalian cells (with respect to the presence of exosome markers and the depletion of non-exosome proteins) to those generated using ultracentrifugation(Skottvoll *et al*., 2019). One part of the reagent and two parts of supernatant were mixed and incubated at 4°C overnight. Pellets of EVs were collected after centrifugation at 10,000 x *g* for 60 minutes at 4°C. The EV pellets were resuspended in PBS for Western blot analysis. For the EV targeting experiments between S2 cells (Fig. 3f), supernatants from transiently transfected donor cells were collected and filtered using 0.22 um PVDF membrane before resuspension and incubation with the recipient cells. Two days before the supernatant swap between EV donor and recipient cell cultures, the recipient cells were transiently transfected with DNA constructs. The recipient cells were incubated in the supernatants with EVs from donor cells for 48 hours at 22°C. For Western blot analyses, cell lysates were prepared using RIPA cell lysis buffer. To measure the size and number of EV particles from S2 cell culture, collected EV pellets were subjected to NTA by System Biosciences, LLC (Palo Alto, CA, USA) (Supp. Fig. 2d-e). The NTA measurements rely on light scattering to extract particle size and the number of particles in a sample and the NTA software (Version 2.3) collects data on multiple particles to calculate the hydrodynamic diameter of each particle using the Stokes-Einstein equation (System Biosciences, LLC).

### Mass spectrometry analysis

Samples were lyophilized and proteins were trypsin-digested as previously described(Pierce et al., 2013). 200ng of digested peptides were analyzed as previously described(Sung et al., 2016). Briefly, peptides were loaded onto a 26-cm analytical HPLC column (75 μm inner diameter) packed with ReproSil-Pur C_18AQ_ 1.9-μm resin (120-Å pore size; Dr. Maisch, Ammerbuch, Germany). Peptides were separated with a 120-min gradient at a flow rate of 350 nl/min at 50°C (column heater) using the following gradient: 2–6% solvent B (7.5 min), 6–25% B (82.5 min), 25–40% B (30 min), 40–100% B (1 min), and 100% B (9 min), where solvent A was 97.8% H_2_O, 2% ACN, and 0.2% formic acid, and solvent B was 19.8% H_2_O, 80% ACN, and 0.2% formic acid. Samples were analyzed using an EASY-nLC 1000 coupled to an Orbitrap Fusion operated in data-dependent acquisition mode to automatically switch between a full scan (*m/z* = 350–1500) in the Orbitrap at 120,000 resolving power and an MS/MS scan of higher-energy collisional dissociation fragmentation detected in the ion trap (using TopSpeed). The automatic gain control (AGC) targets of the Orbitrap and ion trap were 400,000 and 10,000.

### Mass spectrometry data

Raw data were searched using MaxQuant (version 1.5.3.30)(Cox and Mann, 2008; Wagner et al., 2011) against the Uniprot D melanogaster database. Fragment ion tolerance was 0.5 Da. Precursor mass tolerance was 4.5 ppm after automatic recalibration. Searches were permitted up to two missed tryptic peptide cleavages. Cysteine carbamidomethylation was designated as a fixed modification while Methionine oxidation and N-terminal acetylation were designated as variable modifications. False discovery rates were estimated to be <1% using a target-decoy approach. Complete data are in Supp. Table 1.

### Protein expression and purification

To express and purify Arc proteins in the *E. coli* system, the cDNAs of dArc1, dArc2 and rArc were subcloned into the pGEX-4T-1 vectors together with GST-6xHis-tags and TEV protease cleavage site. Arc proteins were expressed in *E. coli* strain BL21 (DE3) grown in LB broth by induction of log-phase cultures with 1 mM isopropyl-β-D-thiogalactopyranoside (IPTG) and incubated overnight at 23°C. Cells were pelleted and resuspended in B-PER lysis buffer (#78243, Thermo Scientific) before centrifugation to collect cell lysates.

Tagged Arc proteins were pulled down using Ni-NTA resin column and the eluates with GST-6xHis-dArc1 and –rArc proteins were used for peptide binding assays (Fig. 4d). For negative stain EM (Fig. 2 and Supp. Fig. 2), the GST-6xHis-tag was removed by TEV protease (#P8112S, NEB) and dArc1 and dArc2 proteins were further purified by size exclusion chromatography using Superdex S200 16/600 (GE Healthcare Life Sciences).

### Western blotting

Proteins were separated by SDS–PAGE, transferred at 200 mA for 60 minutes to nitrocellulose membranes using a Bio-Rad Wet Tank Blotting System in Tris-Glycine Transfer Buffer with 10% methanol. Blocked membranes were incubated with primary antibodies in 0.5% milk PBS-0.1% Tween for overnight. HRP-conjugated antibodies (anti-V5-HRP (#RV5-45P-Z, ICL), anti-mouse IgG HRP (#sc-516102, Santa Cruz Biotechnology), anti-beta-actin-HRP (#HRP-60008, Proteintech), anti-rabbit IgG HRP (#65-6120, Invitrogen), anti-rat IgG HRP (#35470, Invitrogen), anti-alpha-tubulin-HRP (#HRP-66031, Proteintech), anti-cMyc-HRP (#RMYC-45P-Z, ICL), and anti-GST-HRP (#MA4-004-HRP, Invitrogen)) were used at 1:10,000 for 60 minutes. Blots were developed using ECL Western Blotting Substrate (#32109, Pierce), and imaged on a MINI-MED 90 X-Ray Film Processor (AFP Manufacturing Co.).

### Electron microscopy

#### Negative stain EM of purified capsids

dArc1 and dArc2 capsids were examined using negative staining. Briefly, continuous carbon grids (copper, 300 mesh, Electron Microscopy Sciences) were glow discharged for 1 minutes at 15 mA on a PELCO easiGLOW (Ted Pella). 3 uL of sample was applied to grids and allowed to incubate for 60s. Grids were then blotted and stained with 2% (w/v) uranyl acetate solution for 30s. After blotting, grids were allowed to dry for at least 1 hour. Grids were imaged on a Tecnai T12 transmission electron microscope (Thermo Fisher Scientific) operating at 120 kV. Images were recorded on the Gatan Ultrascan camera (Gatan / Ametec).

#### Electron Tomography and Immuno-EM

For imaging of EVs by electron tomography (ET), EVs were prepared as described above. Supernatant was removed and replaced with ∼10 ml 10% Ficoll, 5% sucrose in 0.1M sodium cacodylate trihydrate with minimal disturbance of the pellet. Pellets were transferred to brass planchettes (type A/B; Ted Pella, Inc.) and ultra-rapidly frozen with a HPM-010 high-pressure freezing machine (Bal-Tec/ABRA). Vitrified samples were transferred under liquid nitrogen to cryo-tubes (nunc) containing a frozen solution of 2.5% osmium tetroxide, 0.05% uranyl acetate in acetone and placed in an AFS-2 Freeze-Substitution Machine (Leica Microsystems, Vienna). Samples were freeze-substituted at -90°C for 72 h, warmed to -20°C over 12 h, held at -20° for 12 h, then warmed to room temperature. Samples were rinsed 3x with acetone and infiltrated into Epon-Araldite resin (Electron Microscopy Sciences). Resin was polymerized at 60°C for 24 h.

Serial semi-thin (170 nm) sections were cut with a UC6 ultramicrotome (Leica Microsystems) using a diamond knife (Diatome Ltd., Switzerland). Sections were collected onto Formar-coated copper/rhodium slot grids (Electron Microscopy Sciences) and stained with 3% uranyl acetate and lead citrate. Colloidal gold particles (10 nm) were placed on both surfaces of the grid to serve as fiducial markers for subsequent image alignment. Grids were placed in a dual-axis tomography holder (Model 2040; Fischione Instruments, Inc.) and imaged with a Tecnai T12 transmission electron microscope (Thermo-Fisher Scientific) at 120k eV. For dual-axis tomography, grids were tilted +/- 62° and images acquired at 1° intervals. The grid was rotated 90° and a similar tilt-series was recorded about the orthogonal axis. Tilt-series data was acquired automatically using the SerialEM software package. Tomographic data was calculated, analyzed and modeled on iMac Pro and M1 computers (Apple, Inc) using the IMOD software package.

For immuno-EM, EV pellets were prepared as per above. Supernatant was removed and pellets fixed with 4% paraformaldehyde in PBS for 1 hr. Pellets were then infiltrated with 2.1M sucrose in PBS over 24 h, with >3 changes of the infiltration solution during that time. Pellets were placed onto aluminum sectioning stubs, drained of excess liquid and frozen in liquid nitrogen. Cryosections (100 nm) were cut at -140°C with a UC6/FC6 cryoultramicrotome (Leica Microsystems) using cryo-diamond knives (Diatome Ltd). Cryosections were collected with a wire loop containing 2.3 M sucrose in PBS and transferred to Formvar-coated, carbon-coated, glow-discharged 100-mesh copper/rhodium grids (Electron Microscopy Sciences) at room temperature. Nonspecific antibody binding sites were blocked by incubating the grids with 10% calf serum in PBS for 30’. Sections were then labeled with 1° antibodies (diluted in 5% calf serum/PBS) for 2 h, rinsed 4x with PBS, then labeled with 10 nm and/or 15 nm gold-conjugated 2° antibodies (diluted in 5% calf serum/PBS) for 2 hrs. Grids were rinsed 4x with PBS, 3x with dH_2_O then simultaneously negatively-stained and stabilized with 1% uranyl acetate, 1% methylcellulose in dH_2_O. Immuno-EM samples were imaged as per the tomography samples, above.

#### Immunoprecipitation

For the Myc-co-IP assay (Fig. 4c), transiently transfected S2 cells using Effectene (Qiagen) were cultured in Schneider’s medium at 22 °C for four days. Tagged expression constructs (V5-mCD8^ECD^-sas^TM-ICD^, V5-mCD8^ECD^-APP^ICD^, V5-mCD8^ECD^-Appl^ICD^, dArc1-Myc and Myc-rArc) were cloned in pAc5.1B vector according to standard cloning procedure. For IP analysis, cell lysates were prepared using IP Lysis buffer (#87787, Pierce) and the lysates were incubated in Myc-Trap agarose (#yta-20, Chromotek) following the manufacturer’s protocol and the eluates were analyzed by standard Western blot analysis.

#### Peptide binding assay

For the peptide binding assay (Fig. 4d), biotinylated peptides (wt Sas^ICD^, Sas^ICD^ variations (scrambled and 𝜟YDNPSY), APP^ICD^ and Appl^ICD^) made by RS Synthesis, Inc., were incubated with Streptavidin magnetic beads (#88817, Pierce) for 45 minutes at 4°C and the beads were extensively washed with TBST. Purified GST-6xHis-dArc1 and rArc proteins were added to the beads with bound biotinylated peptides and incubated at 4°C overnight. Similar experiments were performed with Numb PTB domain protein purified from *E. coli*. The beads were carefully washed with TBST and eluates prepared for Western blot analysis following the standard protocol described above.

#### Fluorescent in situ hybridization (FISH)

The FISH protocol was a modification of protocols from (Kosman et al., 2004). Fixed L16 whole embryos were prepared using standard protocols and rinsed with ethanol quickly four times. Then the embryos were permeabilized twice with a mixture of xylenes and ethanol (1:2, v/v) and washed three times with ethanol for 5 minutes each. To rehydrate the embryos, the embryos were washed with 100%, 50% and 0% methanol in PBT sequentially for 30 minutes each step. The rehydrated embryos were permeabilized again using proteinase K (20ug/mL in PBT) for exactly 7 minutes and washed three times for 5 minutes each in PBT followed by a second fixation (5% paraformaldyhyde and 1% DMSO in PBT) for 25 minutes and washed three times in PBT for 5 minutes each. Then the embryos were prepared for pre-hybridization by incubation in 50% hybridization buffer (50% formamide, 5x SSC, 100 μg/ml fragmented salmon testes DNA, 50 μg/ml heparin, 0.1% Tween-20) in PBT for 5 minutes. For pre-hybridization, embryos were incubated in hybridization buffer for more than 90 minutes at 55°C while changing the buffer every 30 minutes. The pre-hybridized embryos were incubated in DIG-tagged dArc1 mRNA probe for 18 hours at 55°C for annealing. The embryos were washed with hybridization buffer three times for 30 minutes each at 55°C, after which the buffer was replaced with replaced the buffer with PBT containing rhodamine-conjugated sheep anti-DIG antibody (#11207750910, SigmaAldrich) overnight at 4°C. Then the embryos were washed and mounted for confocal microscopy.

#### Probe preparation

Probes were designed against a 760 nt region of dArc1 mRNA 3’ UTR sequence, which was used for FISH in a previous study (Ashley *et al*., 2018). To generate antisense and sense probes for dArc1 mRNA, cDNA sequences from *dArc1* were PCR amplified and purified to use as positive and negative probe templates. The DNA templates were heated to 55°C for two minutes and then put back on ice. Transcription reactions were set up to label probes with digoxigenin (DIG, # 11277073910, Roche) and incubated at 37°C for two hours. Probes were precipitated and resuspended in hybridization buffer and stored at -20°C.

The following primers were used to generate dArc1 mRNA probes:

dArc1 probe forward primer: GATTTTTCGTCTGATCCTGGTC

dArc1 probe reverse primer: CCGTTTCTGAGTTTAATGGTTG

## Acknowledgments

Mass spectrometry work was performed at the Caltech Proteome Exploration Laboratory. Imaging was done at the Caltech Biological Imaging facility. EM work was done at the Caltech Cryo-EM facility. We thank Andre Malyutin for negative stain EM. We thank Violana Nesterova for figure preparation. We thank the following colleagues for reagents and *Drosophila* lines: Jason Shepherd (University of Utah) for pGEX-dArc and rArc constructs; Travis Thomson and Vivian Budnik (University of Massachusetts) for rabbit anti-Arc1; Douglas Cavener (Penn State) for rabbit anti-Sas^FL^; Deborah Andrew (Johns Hopkins) for *Sage-GAL4*; James Skeath (Washington University) for guinea pig anti-Numb; Swati Banerjee (UTHSC, San Antonio) for rat anti-Repo, and Yuh-Nung Jan (UCSF) for *UAS-Numb*. We thank Simon Erlendsson, Fernando Bazan, Paul Worley, and Tino Pleiner for discussions about Arc purification and Arc and Sas structures. This work was supported by NIH RO1 grants NS28182 and NS096509 to K.Z., and by Howard Hughes Medical Institute support to R. Deshaies, who was J.M.R.’s faculty supervisor when he was a postdoctoral fellow at Caltech.

## Author Contributions

P. H. L. designed and performed the majority of the experiments. M.A. helped with protein biochemistry work. M.S.L. performed the immuno-EM and EM tomography experiments. J.M.R. performed the mass spectrometry analysis of V5 IPs from EVs. P. H. L. and K.Z. wrote the manuscript. K.Z. directed the project.

## List of Abbreviations

aa: amino acid
AD: Alzheimer’s disease
Ap: apterous
APP: amyloid precursor protein
Co-IP: coimmunoprecipitation
CSP: cell surface protein
ECD: extracellular domain
EM: electron microscopy
EV: extracellular vesicle
EVT: 345 aa region absent from Sas PA/PC isoform
FISH: fluorescence in situ hybridization
FN-III: Fibronectin Type III
GOF: gain of function (overexpression)
ICD: cytoplasmic domain
immuno-EM: immuno-electron microscopy
MVB: multivesicular body
NMJ: neuromuscular junction
NTA: nanoparticle tracking analysis
ORF: open reading frame
PTB: phosphotyrosine binding
Ptp10D: Drosophila receptor tyrosine phosphatase gene located at 10D on the chromosome map
RPTP: receptor tyrosine phosphatase
Sas: Stranded at second
Sas^FL^: full-length Sas (PB/PD isoform)
Sas^short^: Sas isoform lacking EVT region (PA/PC isoform)
SG: salivary gland
TM: transmembrane
UAS: upstream activation sequence
UTR: untranslated region
VFWC: von Willebrand factor C
VNC: ventral nerve cord

## Notes

### Competing Interest Statement

Justin M. Reitsma is affiliated with AbbVie. The author has no other competing interests to declare.

